# Oxytocin receptors in the nucleus accumbens shell are necessary for the onset of maternal behavior in post-parturient mice

**DOI:** 10.1101/2020.11.20.392027

**Authors:** Shannah Witchey, Heather K. Caldwell

## Abstract

Oxytocin (Oxt) signaling via its receptor, the Oxt receptor (Oxtr), is important to the onset of mammalian maternal care. Specifically, evidence suggests that Oxt signaling around the time of parturition underlies the critical shift in how pups are perceived, i.e. from aversive stimuli to rewarding stimuli. Previous work from our lab has found that both Oxtr knockout (−/−) mice and forebrain-specific Oxtr knockout (FB/FB) are more likely than controls to abandon their first litters. Based on these data, we hypothesized that this observed pup abandonment phenotype was due to a failure of the brain to “switch” to a more maternal state. In order to identify where in the brain Oxt signaling contributes to the onset of maternal care we performed three experiments. In Experiment 1, virgin Oxtr FB/FB females were assessed for genotypic differences in maternal behavior and c-Fos expression following maternal sensitization was quantified. In Experiment 2, c-Fos expression was quantified in Oxtr −/− and Oxtr FB/FB females following parturition. In Experiment 3, based on our findings from Experiment 2, the Oxtr in the nucleus accumbens shell (NAcc) was genetically deleted in female Oxtr floxed mice (Oxtr Flox/Flox) mice using a Cre recombinase expressing adeno-associated virus. In Experiment 1, sensitized virgin Oxtr FB/FB females had significantly lower retrieval latencies on the first day of testing and reduced c-Fos expression in the dorsal lateral septum compared to controls. In Experiment 2, increased c-Fos expression was observed in the NAcc shell of both Oxtr −/− and Oxtr FB/FB dams as compared to controls. In Experiment 3, virally mediated knockout of the Oxtr in the NAcc shell completely disrupted the onset of maternal care. Thus, by genetically deleting Oxtr expression in the NAcc the pup abandonment phenotype previously observed in Oxtr −/− and Oxtr FB/FB dams was recreated. Taken together, these data suggest that in post-parturient mice, Oxtr expression in the NAcc shell is critical to the onset of maternal behavior.

## Introduction

In mammals, the neuropeptide oxytocin (Oxt), known for its role in the facilitation of uterine contractions and milk ejection, acts within the central nervous to help initiate maternal behaviors (1-3). In the peripartum period there are marked increases in the peripheral release of Oxt as well as increases in Oxt within the paraventricular (PVN) and supraoptic (SON) nuclei of the hypothalamus and limbic regions of the brain (4, 5). During late pregnancy and at parturition there is a rapid upregulation, and subsequent down regulation, of the Oxt receptor (Oxtr), which is the single identified receptor subtype for Oxt. In rodents, Oxtr expression, within numerous brain regions, is upregulated by gestational day 22, maintained at parturition, and downregulated in the post-partum period (6-8). Perhaps not surprisingly, the pattern of Oxtr upregulation in specific brain regions differs between parturient and pup-sensitized virgin rats within several brain regions, including the bed nucleus stria terminalis (BNST), the medial preoptic area (mPOA), and the medial amygdala (MeA) (6, 7). However, while parturient and pup-sensitized virgin females differ in their Oxtr expression, both have similar increases in peripheral Oxt concentrations (9). Thus, there are significant, and measurable, differences in Oxt neurochemistry in a mother who has given birth as compared to one that has not.

Data also suggest that Oxtr expression is precisely regulated around the time of birth. Specifically, temporally controlled Oxtr expression appears to be the greatest in brain regions that have been identified as part of the maternal behavior neural network (MBNN). The MBNN includes key neural sites where sensory cues and cortical inputs, as well as hormones and neurochemicals, coordinate and modulate the expression of maternal care (10, 11). In the MBNN, the mPOA and BNST integrate pup-related sensory input and relay that information to the ventral tegmental area (VTA). The VTA then dampens nucleus accumbens (NAcc) output causing a disinhibition of the ventral pallidum (VP), which is permissive for the triggering of maternal responsiveness (12, 13). However, as mentioned above, the way that pup-related sensory input is interpreted by the brain does differ in parturient versus virgin pup-sensitized dams. Specifically, in mice, immediate early gene studies, e.g. c-Fos, suggest that there is differential neuronal activation within the BNST, the mPOA, the VTA, the NAcc core, and the NAcc shell in parturient versus virgin pup-sensitized dams following exposure to pups (14). Perhaps most importantly though, there is evidence that the Oxt system directly mediates aspects of maternal behavior.

Pharmacological studies have determined that Oxt acting via the Oxtr is particularly important for the onset of maternal behavior. In gonadal steroid-primed virgin female rats Oxt induces (15) and antagonism of the Oxtr impairs the onset of maternal care, which suggests that that Oxt plays a critical role in pup-sensitization (16, 17). To date, two brain regions have been identified that are thought to be critical for Oxt’s effects on the onset of maternal behavior, the mPOA and the VTA. The mPOA has estrogen-mediated increases in Oxtr expression (7) that increase its sensitivity to pup stimuli. The VTA also expresses the Oxtr, specifically on dopamine and glutamate neurons (18), and it is here that Oxt signaling through its receptor resulted in increases in dopamine signaling in the NAcc, ultimately contributing to the onset of maternal care as well as individual differences in licking and grooming (7, 19). The timing of the Oxt signal is also important, as an Oxtr antagonist (Oxtr-A) administered at the time of parturition can block the onset of maternal behavior without having long-term effects on established maternal care (20). This is consistent with evidence suggesting that once maternal behavior is established the administration of Oxt cannot enhance it further. Though, it is of note that administration of an Oxtr-A can reduce pup-directed behaviors (16, 21). Taken together these data suggest that Oxt signaling through the Oxtr is important in the initiation, and in specific instances the maintenance, of pup-directed maternal responses.

One way that the contributions of Oxt to maternal care have been interrogated is through the use of mice with genetic disruptions of their Oxt system. Specifically, Oxt knockout (Oxt −/−) and Oxtr knockout (Oxtr −/−) mice. Interestingly, Oxt −/− and Oxtr −/− mice have no deficits in fertility, pregnancy, parturition, or maternal care (if it is initiated). Though, in both lines, due to their inability to milk eject, and thus nurse their offspring, there is complete litter loss within 24 hours of parturition unless the pups are cross fostered (22, 23). That said, these mice do have issues with the initiation of maternal care. In a study from our lab, which separated nursing from maternal care, we found that if an Oxtr −/− dam initiated maternal care the quality of maternal care is not compromised. However, there is a robust pup abandonment phenotype, with 67% of Oxtr −/− dams abandoning their pups as compared to only 20% of Oxtr +/+ dams (24). A similar phenotype is observed in Oxtr forebrain conditional knockout (Oxtr FB/FB) dams, which are able to nurse their young. In these mice, 40% of dams abandon their first litter as compared to 10% in controls (25). The data with respect to maternal sensitization in Oxt −/− virgin females are conflicting, with one study reporting deficits in retrieval latencies and pup licking (26) and another reporting normal sensitization to pups (23). Oxtr −/− dams are known to spend equal amounts of time crouching over their pups as Oxtr +/+ dams; though, in Oxtr −/− dam’s cages, pups are found to be scattered in the nest more frequently within the first 24 hours (23). These data highlight the importance of Oxtr signaling for the onset of maternal care, as well as how a sensitized female’s Oxt system likely differs from that of a post-parturient female (27-33).

Pharmacological and genetic animal models have clearly demonstrated that Oxt is important for the onset of maternal care. Here we propose that another brain region, or node, for Oxt signaling is also critical for the onset of maternal care following parturition. Given the aforementioned pup abandonment phenotype observed in Oxtr −/− and Oxtr FB/FB dams (24, 25), our laboratory has been particularly interested in identifying the neural substrates where Oxt acting via the Oxtr affects the onset of maternal care in post-parturient mice. We *hypothesized* that our observed pup abandonment phenotype was due to a failure of the brain to “switch” to a more maternal state. We predicted that Oxt acting via the Oxtr in brain regions important to the reward system, (e.g. VTA, NAcc, and VP) were necessary for a shift in the perception of pups from aversive stimuli to rewarding stimuli (34).

To test this hypothesis, we examined neuronal activation by quantifying immediate early gene induction, i.e. c-Fos, in sensitized Oxtr FB/FB dams (Experiment 1) as well as in Oxtr −/− and Oxtr FB/FB dams one-hour following parturition (Experiment 2). Based on our findings from Experiment 2 we then went on to genetically disrupt the Oxtr in a brain region of interest to evaluate its contributions to the onset of maternal care (Experiment 3). Taken together, these studies sought to identify brain regions that differ in their Oxt signaling-dependent activation between post-parturient and sensitized females as well as determine where in the brain the Oxt system signals to facilitate the onset of maternal care.

## Results

### Experiment 1: Maternal sensitization and immediate early gene activation in Oxtr +/+ and Oxtr FB/FB virgin, naïve females

A repeated measures ANOVA revealed a main effect of day of testing for retrieving the first pup (F_1,12_=8.827, p=0.012) and retrieving all pups (F_1,11_=51.046, p=0.005) (Figure 1). (One animal was not included in the statistical analysis for the retrieval of all pups due to their only having two foster pups compared to rest of the females having four foster pups). Additionally, an interaction of day and genotype was found for retrieval of first pup (F_1,12_=5.036, p=0.044) and retrieval of all pups (F_1,11_=15.550, p=0.002). Post hoc analysis revealed a genotypic difference in the latency to retrieve the first pup (F_1,13_=6.038, p=0.030) and all pups (F_1,13_=10.775, p=0.015) on the first day of exposure, with the Oxtr FB/FB females having lower retrieval latencies compared to Oxtr +/+ females. No genotypic differences in latencies to retrieve first pup were observed on Day 2 (F_1,13_=0.666, p=0.430), Day 3 (F_1,13_=0.678, p=0.426) or Day 4 (F_1,13_=0.548, p=0.548). Additionally, no differences were observed between Oxtr +/+ or Oxtr FB/FB females in the latency to retrieve all pups on Day 2 (F_1,13_=015, p=0.906), Day 3 (F_1,13_=0.173, p=0.686), or Day 4 (F_1,13_=1.080, p=0.323).

**Figure 1:**
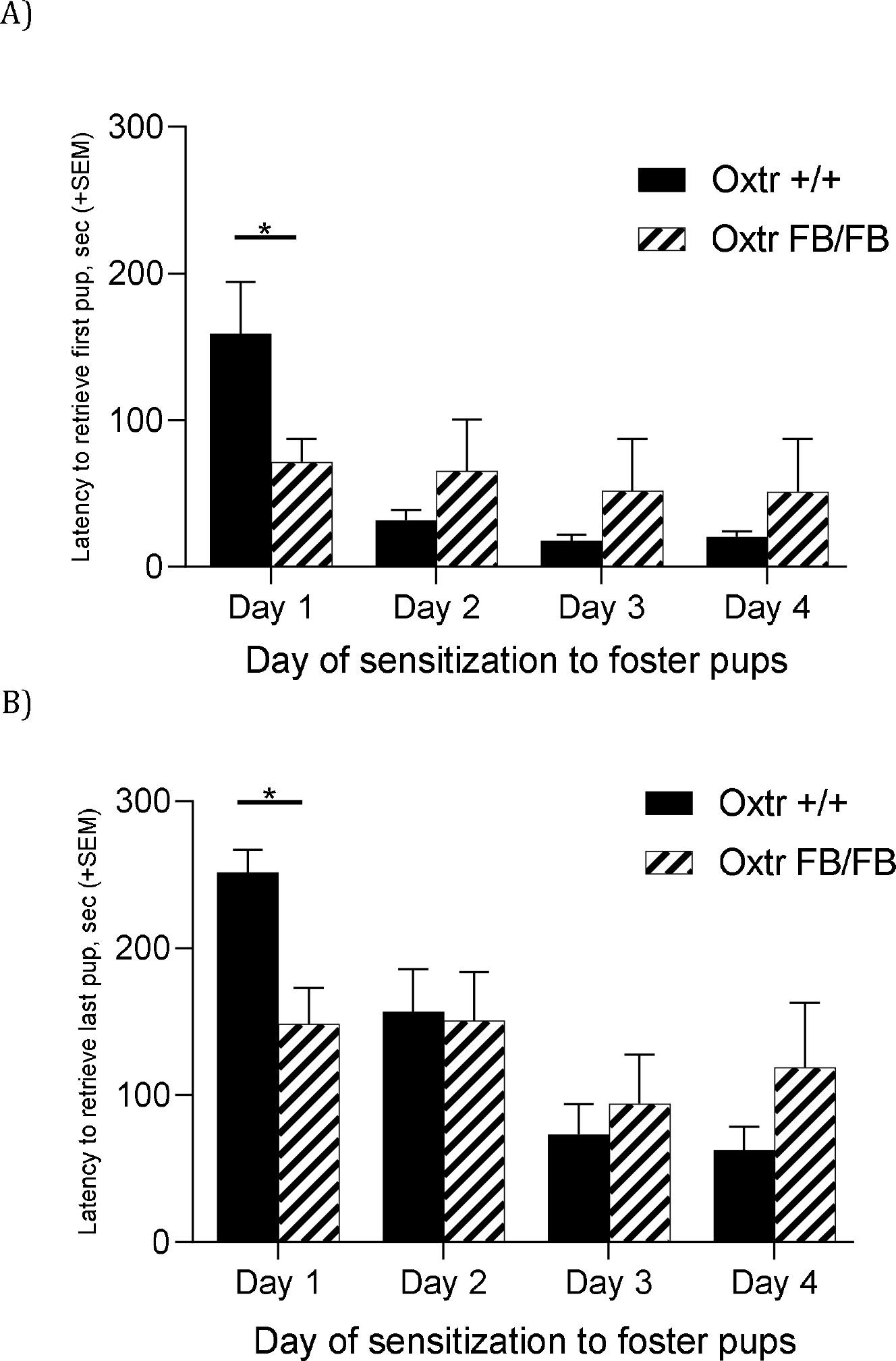
Graphs of latency to retrieve (A) first pup or (B) all pups across four days of sensitization in wildtype (Oxtr +/+) and Oxtr conditional forebrain knockout (Oxtr FB/FB) naïve, virgin females. On day one, Oxtr FB/FB females displayed decrease latency to retrieve first and all pups. Graphs depict mean + SEM (*p≤0.05).

Sensitized Oxtr FB/FB (n=6) and Oxtr +/+ (n=8) females did not differ in any maternal behaviors observed across testing days (Figure 2). No genotypic differences were observed on any of the days. Day 1: duration time on nest (F_1,12_=2.111, p=0.174), nest building (F_1,12_=0.148, p=0.708), self-grooming (F_1,12_=0.021, p=0.887), nonsocial (F_1,12_=0.083, p=0.778) and pup interactions (F_1,12_=0.097, p=0.762). Day 2: duration of time on nest (F_1,12_=0.626, p=0.446), nest building (F_1,12_=0.182, p=0.678), self-grooming (F_1,12_=0.023, p=0.882), nonsocial (F_1,12_=0.012, p=0.915) and pup interactions (F_1,12_=1.330, p=0.273). Day 3: time on nest (F_1,12_=0.318, p=0.584), nest building (F_1,12_=2.757, p=0.125), self-grooming (F_1,12_=0.003, p=0.959), nonsocial (F_1,12_=0.720, p=0.414) and pup interactions (F_1,12_=1.040, p=0.330). Day 4: time on nest (F_1,12_=1.790, p=0.208), nest building (F_1,12_=0.913, p=0.360), self-grooming (F_1,12_=4.486, p=0.058), nonsocial (F_1,12_=1.142, p=0.308) and pup interactions (F_1,12_=0.001, p=0.975).

**Figure 2:**
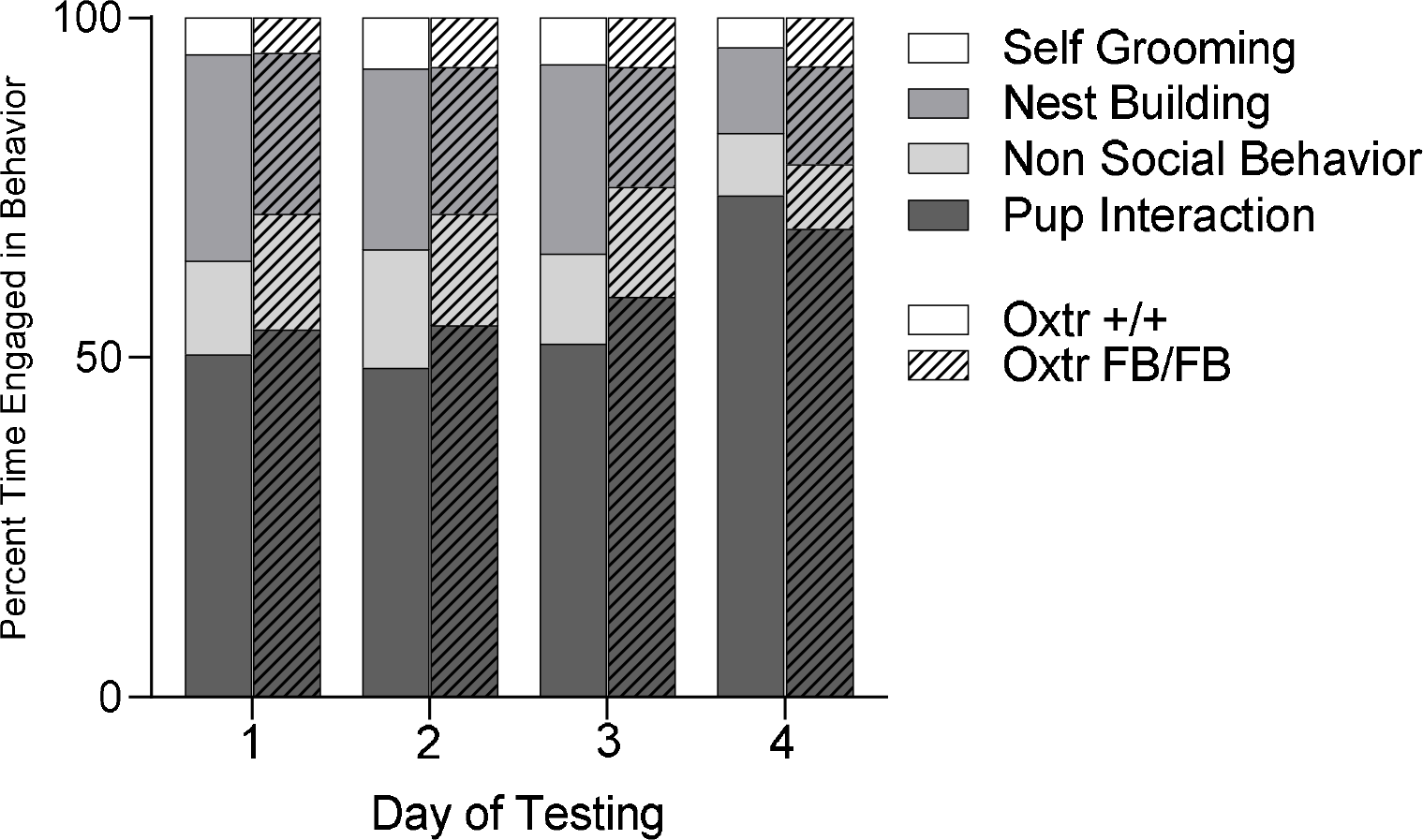
Graph of maternal behaviors observed across the four days of sensitization of wildtype (Oxtr +/+) and Oxtr conditional forebrain knockout (Oxtr FB/FB) virgin, naïve females. Each female was exposed to foster pups for four consecutive days to induce maternal sensitization. Females did not differ in any maternal behaviors observed across testing days and there were no genotypic differences observed on any of the days. The graph depicts percent of time performing behavior.

The Oxtr FB/FB sensitized females had decreased c-Fos immunoreactivity in the LSD compared to the Oxtr +/+ sensitized females (F_1,10_=11.339, p=0.008) (Figure 3 and 4). No genotypic differences were observed in the BNSTD (F_1,10_=0.143,p=0.714), the BNSTV (F_1,10_=0.453, p=0.518), the LHB (F_1,10_=0.001, p=0.973), the LSV (F_1,10_=2.616, p=0.140), the NAcc core (F_1,10_=0.172, p=0.688), the NAcc shell (F_1,10_=0.393, p=0.546), the MeA (F_1,10_=0.185, p=0.678), the mPOA (F_1,10_=1.113, p=0.319), or the PAG (F_1,10_=0.249, p=0.630).

**Figure 3:**
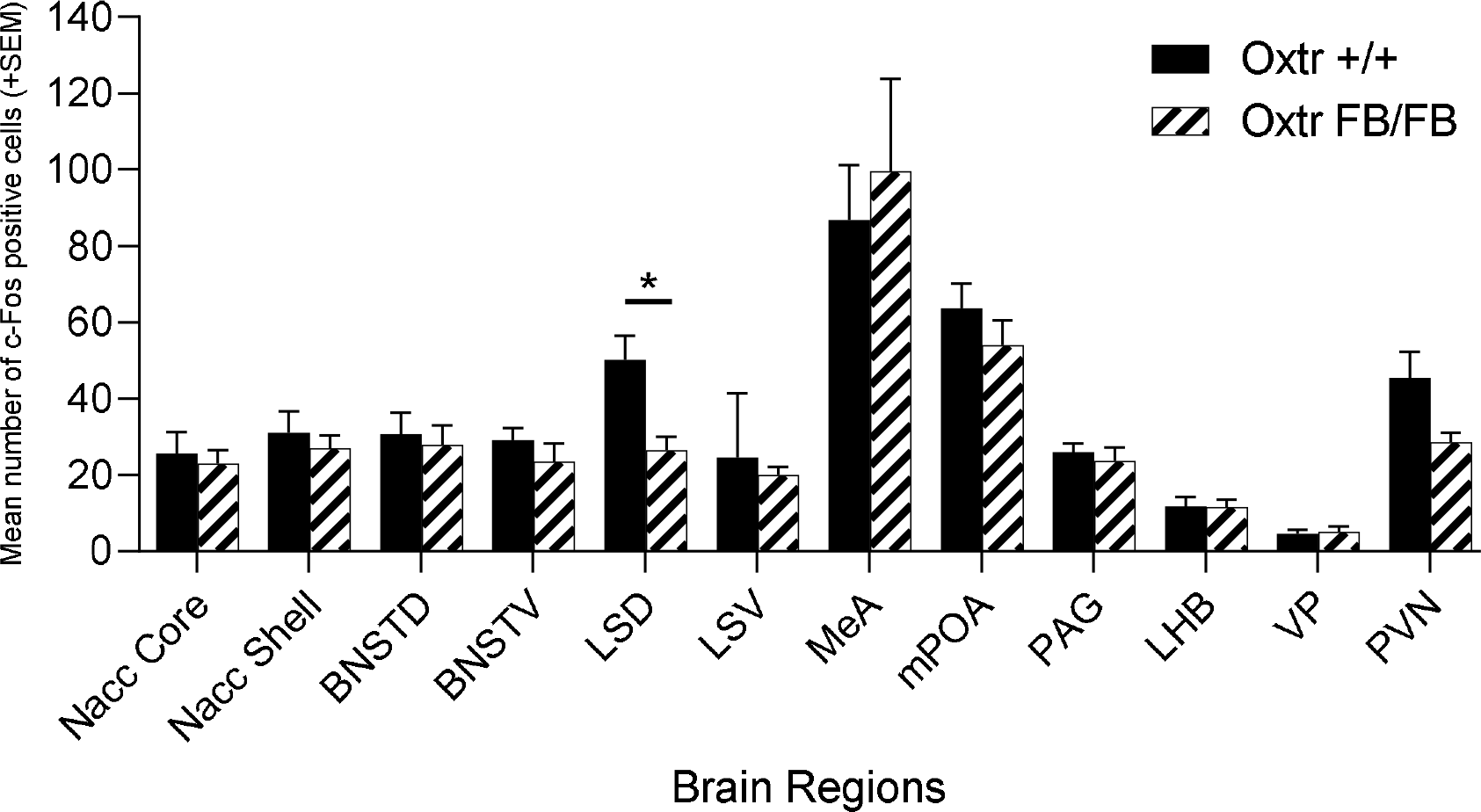
Number of c-Fos immunoreactive cells per brain region in wildtype (Oxtr +/+) and Oxtr conditional forebrain knockout (Oxtr FB/FB) females sensitized to pups. On the fourth day of repeated exposure to pups, the Oxtr +/+ and Oxtr FB/FB have a significant genotypic difference in c-Fos immunoreactivity within the lateral septum. The Oxtr FB/FB females have an increase in activation in the NAcc shell compared to Oxtr +/+ female. No genotypic differences were observed in any of the other measured brain regions. Graphs depict mean + SEM (*p≤0.05). BNSTD, dorsal bed nucleus stria terminalis; BNSTV, ventral bed nucleus stria terminalis; LHB, lateral habenular nuclei; LSD, dorsal lateral septum; LSV, ventral lateral septum; MeA, medial amygdala; mPOA, medial preoptic area; NAccC, nucleus accumbens core; NAccS, nucleus accumbens shell; PAG, periaqueductal grey

**Figure 4:**
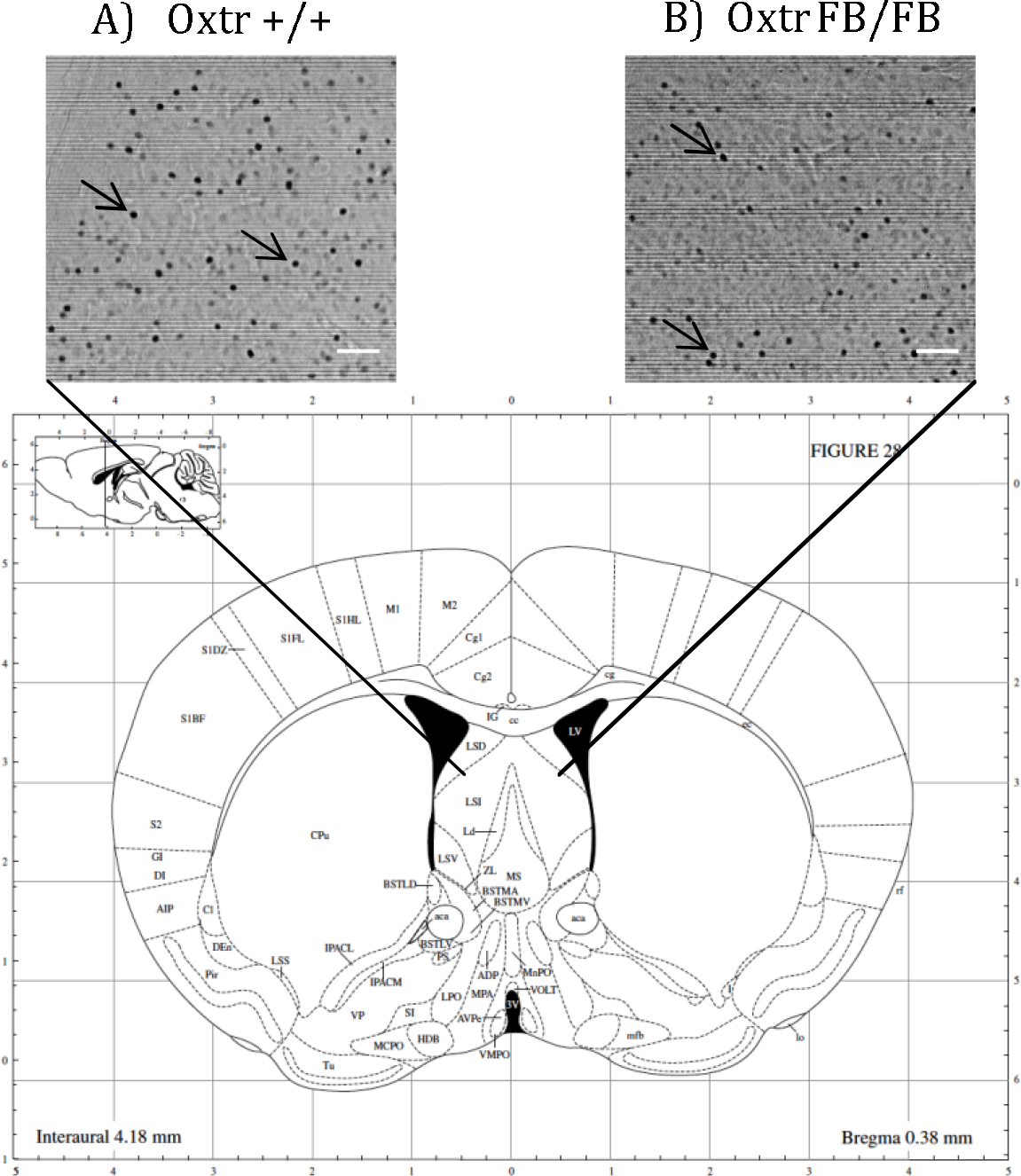
Photomicrograph of the lateral septum (LS) showing c-Fos immunoreactivity in (A) wildtype (Oxtr +/+) and (B) Oxtr conditional forebrain knockout (Oxtr FB/FB) sensitized females one hour after the fourth day of exposure to pups. C) Illustration of the corresponding coronal section of the mouse brain (Paxinos and Franklin 2001). The arrows indicate example of c-Fos immunopositive cell at x100 magnification and the scale bar represents 100 µm.

### Experiment 2: Parturition-induced immediate early gene activation in Oxtr +/+, Oxtr −/−, and Oxtr FB/FB dams

In the one-hour post-parturient Oxtr +/+ (n=8) and Oxtr FB/FB (n=6) dams there was a significant genotypic difference in c-Fos immunoreactivity within the NAcc shell (F_1,13_=5.192, p= 0.042), with an increase in activation in Oxtr FB/FB dams compared to Oxtr +/+ dams (Figures 5 & 6). No genotypic differences in c-Fos activation were observed in any of the other measured brain regions: BNSTD (F_1,13_=0.594, p=0.456), BNSTV (F_1,13_=0.257, p=0.621), LHB (F_1,13_=0.330,p=0.576), LSD (F_1,13_=0.163, p=0.694), LSV (F_1,13_=0.021, p=0.888), NAcc core (F_1,13_=0.130, p=0.725), MeA (F_1,13_=0.036, p=0.852), mPOA (F_1,13_=1.421, p=0.264), PAG (F_1,13_=0.106, p=0.751), PVN (F_1,13_=0.027, p=0.872), VP (F_1,13_=0.608, p=0.451) or VTA (F_1,13_=0.597, p=0.457).

**Figure 5:**
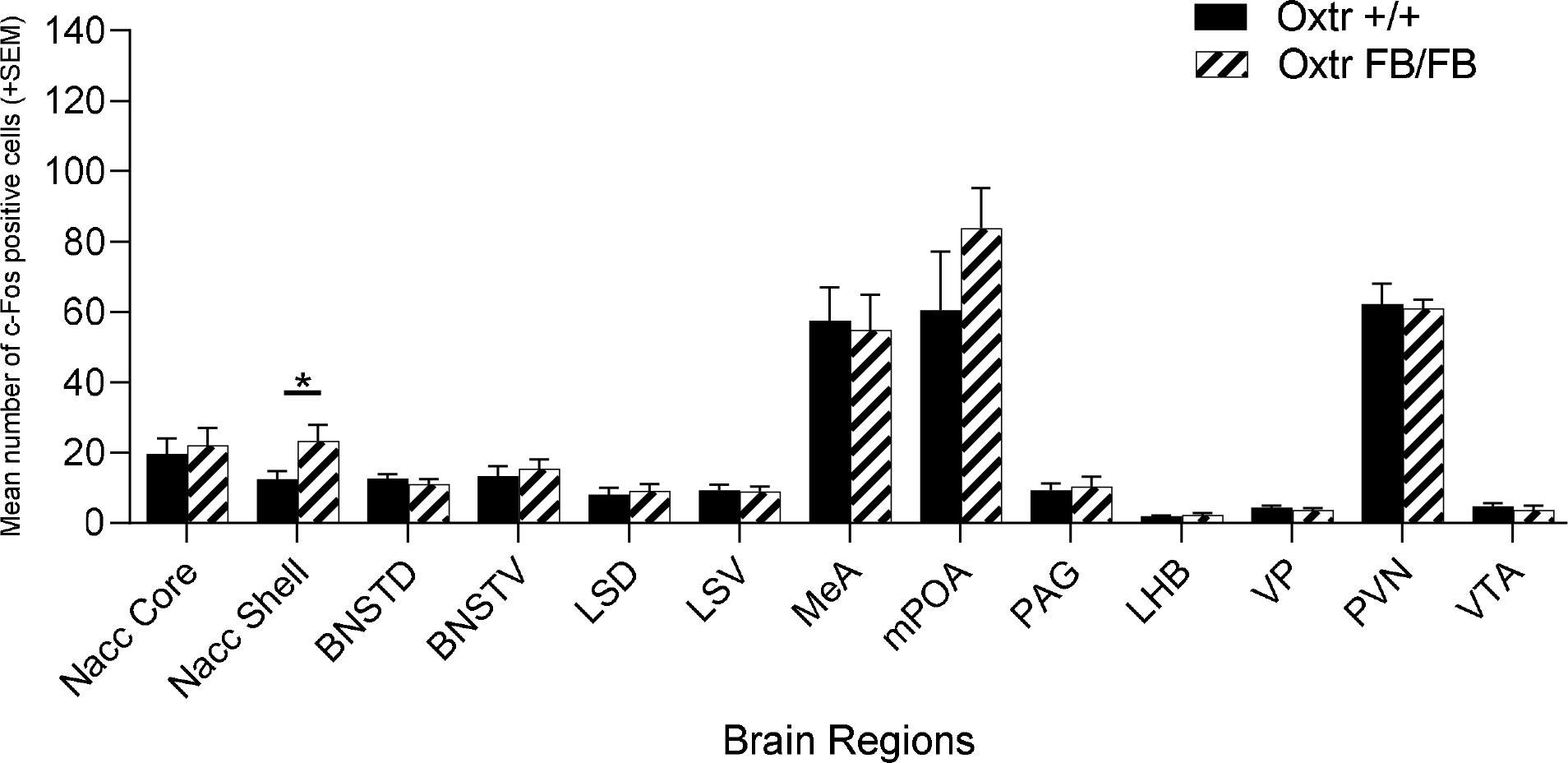
Number of c-Fos immunoreactive cells per brain region in wildtype (Oxtr +/+) and Oxtr conditional forebrain knockout (Oxtr FB/FB) dames following parturition. In the one-hour post parturient Oxtr +/+ and Oxtr FB/FB a significant genotypic difference in c-Fos immunoreactivity was found within the NAcc shell. The Oxtr FB/FB dams have an increase in activation in the NAcc shell compared to Oxtr +/+ dam. No genotypic differences were observed in any of the other measured brain regions. Graphs depict mean + SEM (*p≤0.05). BNSTD, dorsal bed nucleus stria terminalis; BNSTV, ventral bed nucleus stria terminalis; LHB, lateral habenular nuclei; LSD, dorsal lateral septum; LSV, ventral lateral septum; MeA, medial amygdala; mPOA, medial preoptic area; NAcc core, nucleus accumbens core; NAcc shell, nucleus accumbens shell; PAG, periaqueductal grey

**Figure 6:**
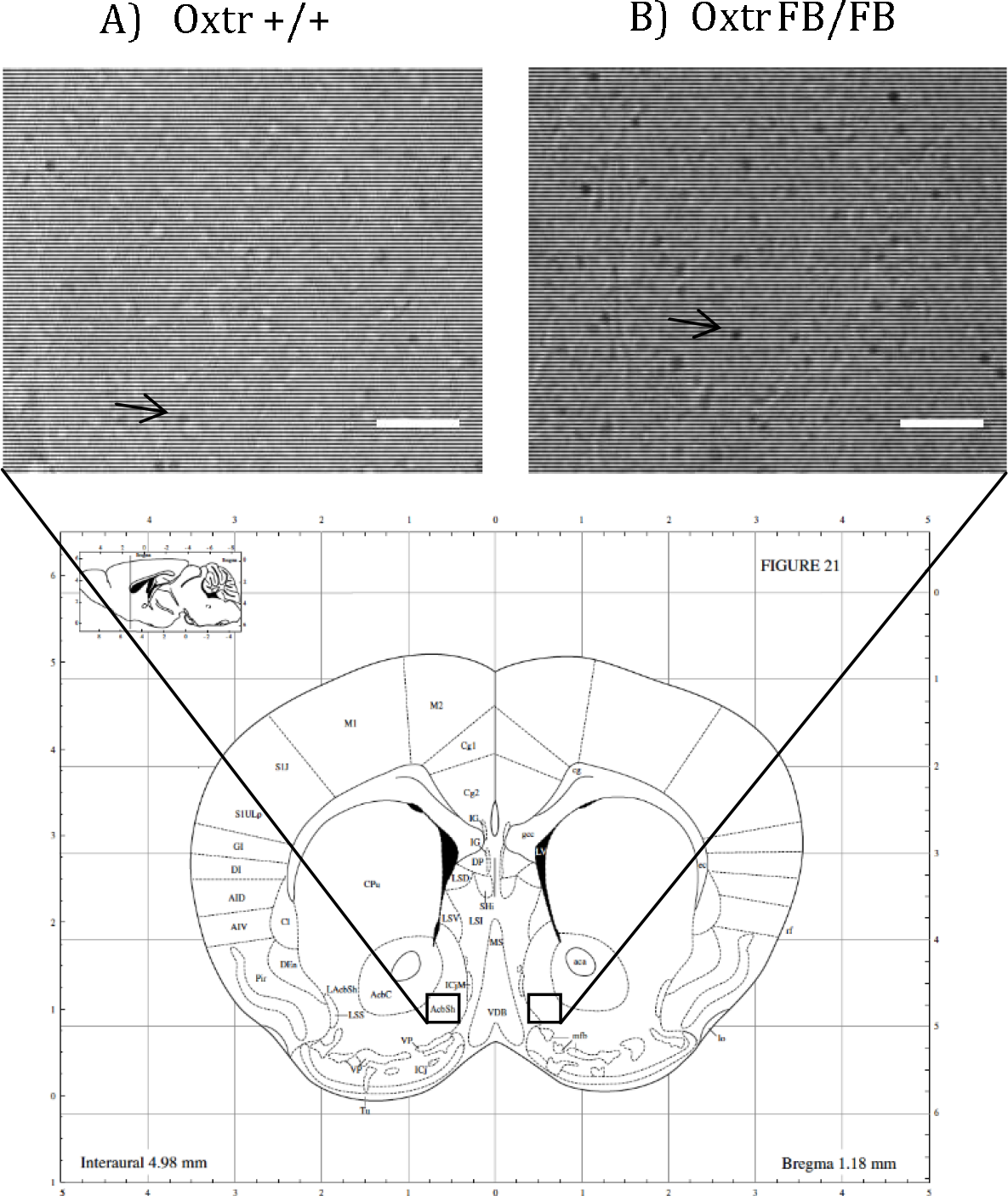
Photomicrographs of the nucleus accumbens (NAcc) shell showing c-Fos immunoreactivity in (A) wildtype (Oxtr +/+) and (B) Oxtr conditional forebrain knockout (Oxtr FB/FB) dams one hour after parturition. C) Illustration of the corresponding coronal section of the mouse brain (Paxinos and Franklin 2001). The arrows indicate example of c-Fos immunopositive cell at x100 magnification and the scale bar represents 100 µm.

In the one-hour post-parturient Oxtr +/+ (n=7) and Oxtr −/− (n=9) dams there was a significant genotypic difference in c-Fos immunoreactivity within the NAcc shell (F_1,15_=12.756, p= 0.03) with an increase in activation in Oxtr FB/FB dams compared to Oxtr +/+ dams (Figures 7 & 8) and the VP (F_1,15_=12.938, p=0.003) with an increase in activation in Oxtr +/+ dams. No genotypic differences in c-Fos expression were observed in any of the other measured brain regions: BNSTD (F_1,15_=1.990, p=0.180), BNSTV (F_1,15_=0.133, p=0.721), LHB (F_1,15_=0.004, p=0.950), LSD (F_1,15_=0.284, p=0.603), LSV (F_1,15_=0.032, p=0.860), MeA (F_1,15_=0.512, p=0.486), mPOA (F_1,15_=0.781, p=0.392), NAcc core (F_1,15_=0.856, p=0.371), PAG (F_1,15_=0.419, p=0.528), PVN (F_1,15_=0.012, p=0.916) or VTA (F_1,15_=3.354, p=0.088).

**Figure 7:**
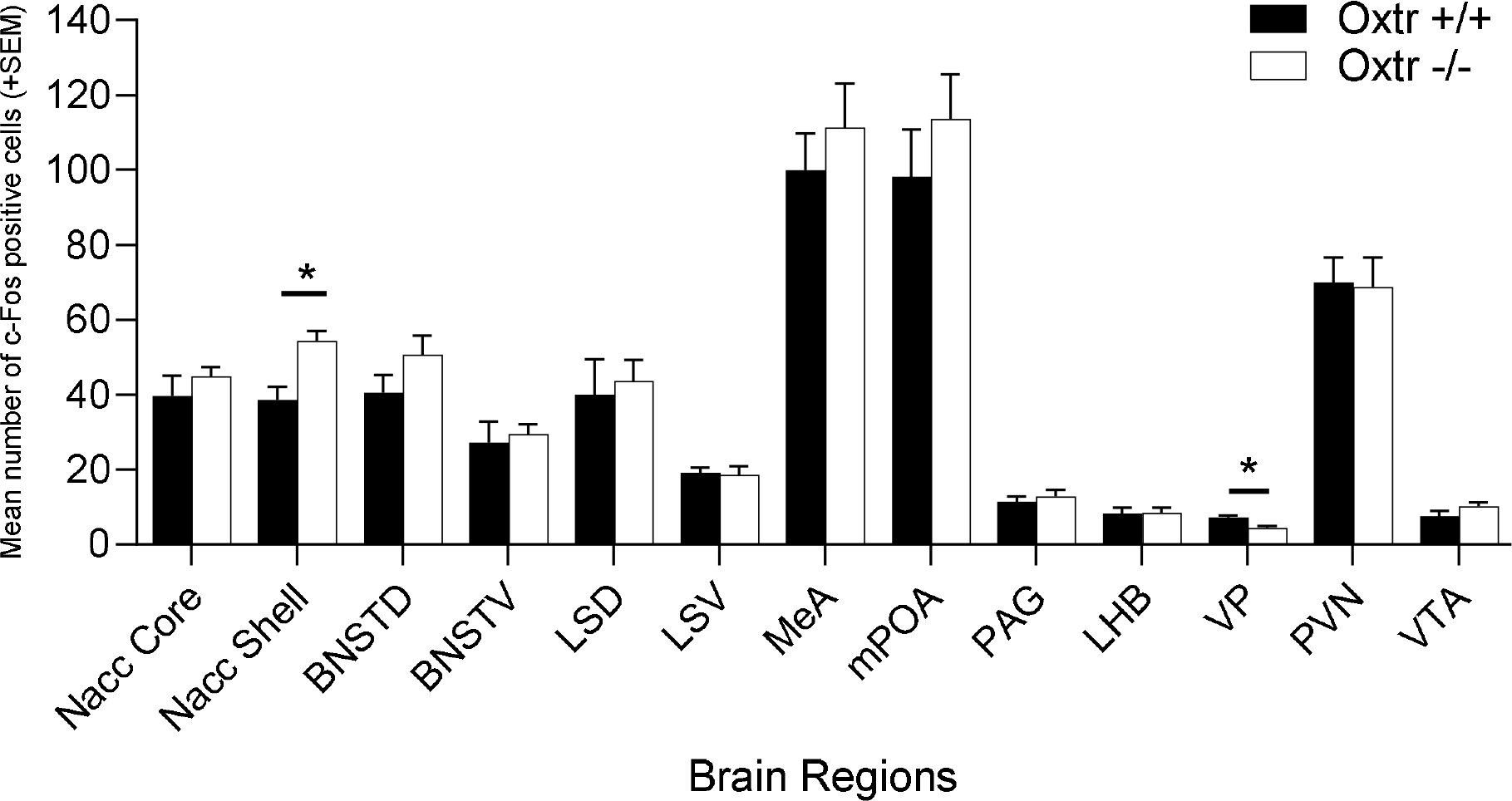
Number of c-Fos immunoreactive cells per brain region in wildtype (Oxtr +/+) and Oxtr knockout (Oxtr −/−) dams following parturition. In the one-hour post parturient Oxtr +/+ and Oxtr −/− a significant genotypic difference in c-Fos immunoreactivity was found within the NAcc shell and VP. The Oxtr −/− dams have an increase in activation in the NAcc shell and decreased activation in the VP compared to Oxtr +/+ dam. No genotypic differences were observed in any of the other measured brain regions. Graphs depict mean + SEM (*p≤0.05). BNSTD, dorsal bed nucleus stria terminalis; BNSTV, ventral bed nucleus stria terminalis; LHB, lateral habenular nuclei; LSD, dorsal lateral septum; LSV, ventral lateral septum; MeA, medial amygdala; mPOA, medial preoptic area; NAcc core, nucleus accumbens core; NAcc shell, nucleus accumbens shell; PAG, periaqueductal grey.

**Figure 8:**
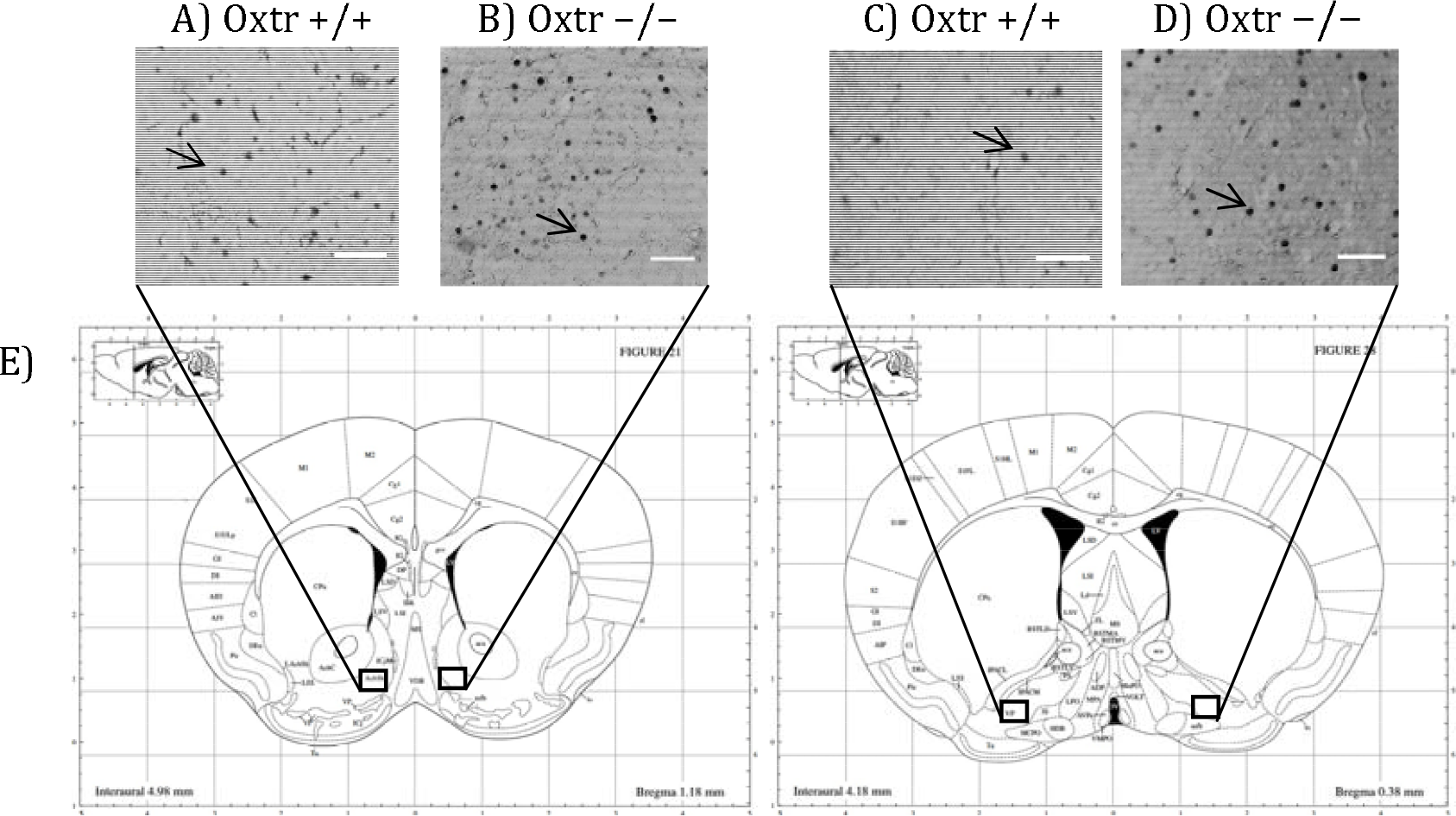
Photomicrographs of the nucleus accumbens (NAcc) shell and ventral pallidum (VP) showing c-Fos immunoreactivity one hour after parturition in in wildtype (Oxtr +/+) and Oxtr knockout (Oxtr −/−) dams. A) and B) are representative images of nucleus accumbens (NAcc) shell. C) and D) are representative images of the ventral pallidum (VP). E) Illustration of the corresponding coronal section of the mouse brain (Paxinos and Franklin 2001). The arrows indicate example of c-Fos immunopositive cell at x100 magnification and the scale bar represents 100 µm.

### Experiment 3: Effects of targeted genetic knockdown of the Oxtr on maternal behavior

In total, n=7 Cre-IRES-GFP and n=8 IRES-GFP female littermates were evaluated in this experiment. Site checks verified that the viral injections were all within the NAcc shell, though a few were prior to or immediately after the targeted bregma position at +1.58 anterior posterior (Figure 9). One animal was removed from the experiment due to disturbances in the animal facility during testing. No notable differences in general health of the dam or pups on PND0 were observed between treatment groups. All pups were found alive on PND0, except for one CRE-IRES-GFP dam. Since the majority of litter losses occurred by PND1, pup weights were only compared on PND0 and no treatment specific differences were observed (F_1,12_=1.114, p=0.287)(Figure 10). A Fisher’s exact test identified a significant genotypic difference in pup abandonment as a function of treatment (F_1,14_=6.583, p=0.023). Approximately, 71% of females injected with Cre-IRES-GFP abandoned their litters by PND2, with all pups cannibalized or found dead in their cage. In contrast, only 12.5% of dams injected with IRES-GFP abandoned their litters by PND2 (Figure 11). This pup abandonment extended to second litters with 40% of Cre-IRES-GFP dams abandoning their litters compared to 0% of IRES-GFP dams. Regardless of litter survival, all dams were tested for anxiety-like behavior in the elevated plus maze and the open field, and for depression-like behavior in the forced swim test. No treatment-specific differences were observed in the percent time spent on the open arms of elevated plus (F_1,14_=0.126, p=0.728), the percent time in the inner arena of open field (F_1,14_=0.337, p=0.572), or percent time spent swimming in the forced swim test (F_1,14_=0.249, p=0.626) (Figure 12). Due to the high percentage of pup abandonment in Cre-IRES-GFP dams, no maternal behaviors were scored.

**Figure 9:**
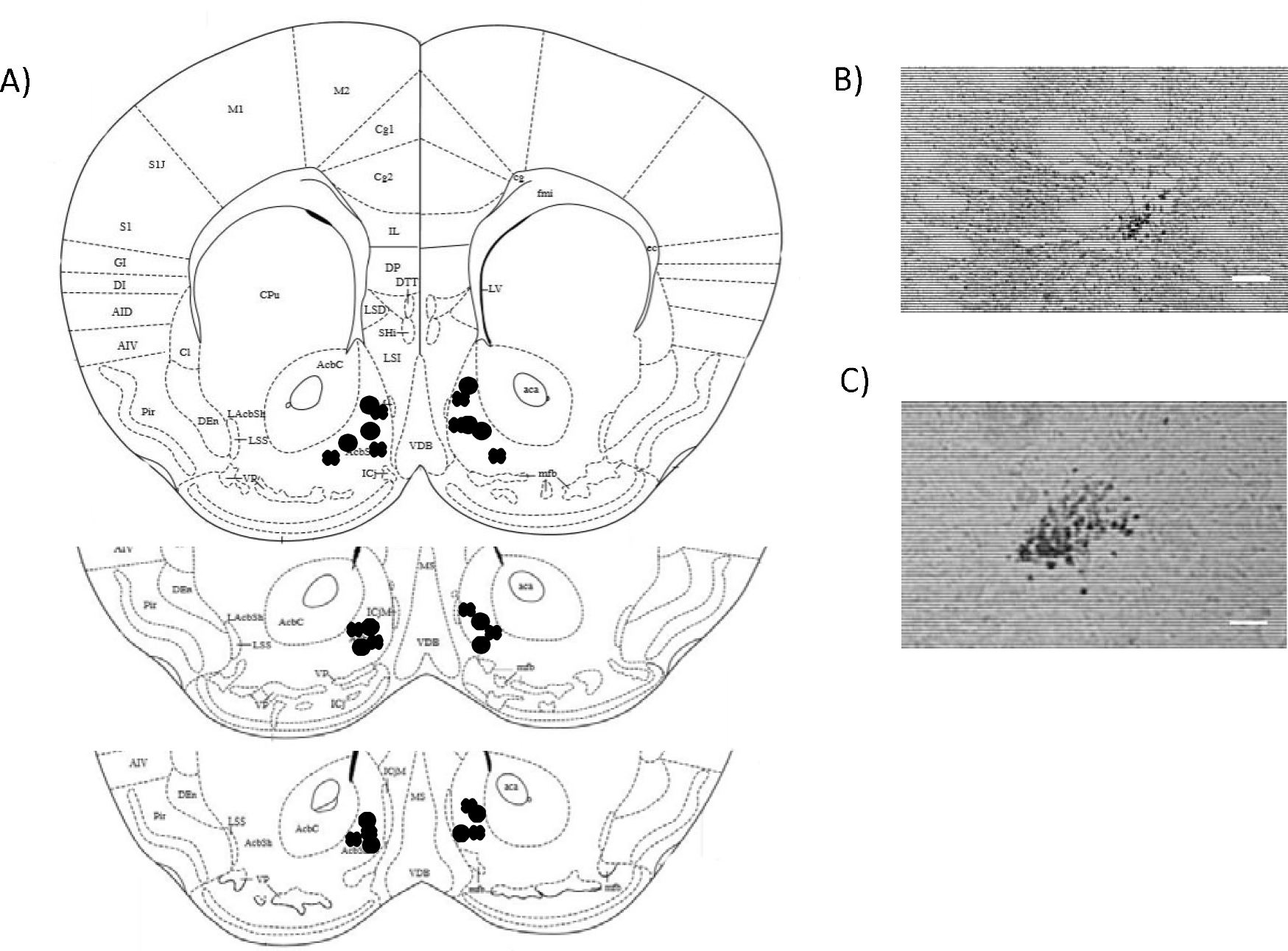
Site checks for Oxtr^flox/flox^ dams receiving viral injections in the nucleus accumbens shell illustrated on corresponding coronal sections of the mouse brain (Paxinos and Franklin 2001). A) The “O”‘s represent injections sites control virus injected dams (IRES-GFP) and “X”‘s represent Cre (IRES-CRE) targeted injections in the nucleus accumbens shell. B) and C) are photomicrographs representatives of GFP labelled cells in IRES-GFP and IRES-CRE, respectively. The scale bar represents 100 µm.

**Figure 10:**
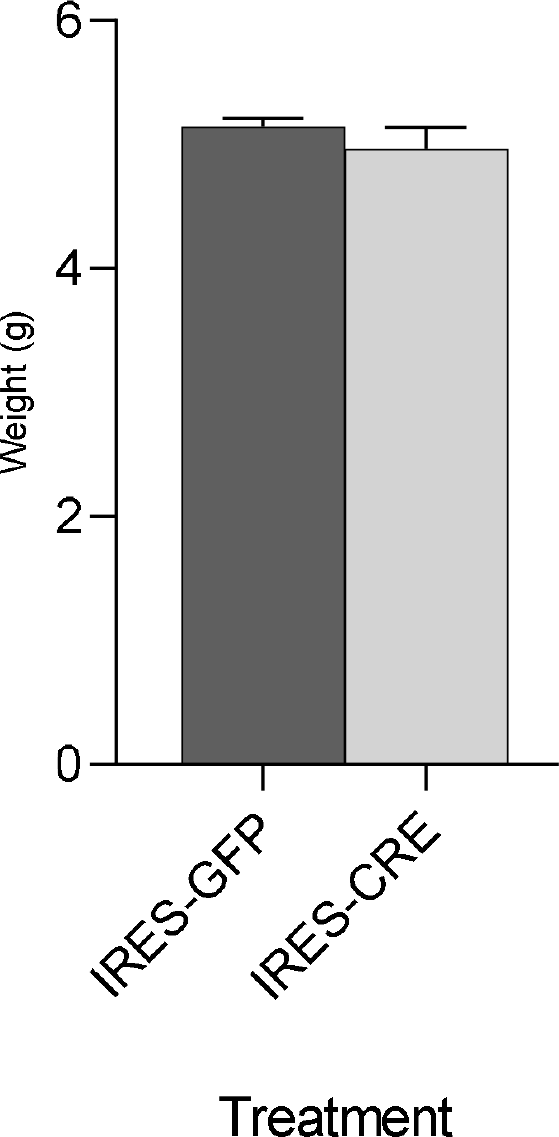
Weights on PND0 of pups from Oxtr^flox/flox^ dams that received either control (IRES-GFP) or Cre (IRES-CRE) targeted injections in the nucleus accumbens (NAcc) shell. No significant genotypic difference in pup weight by treatment was observed.

**Figure 11:**
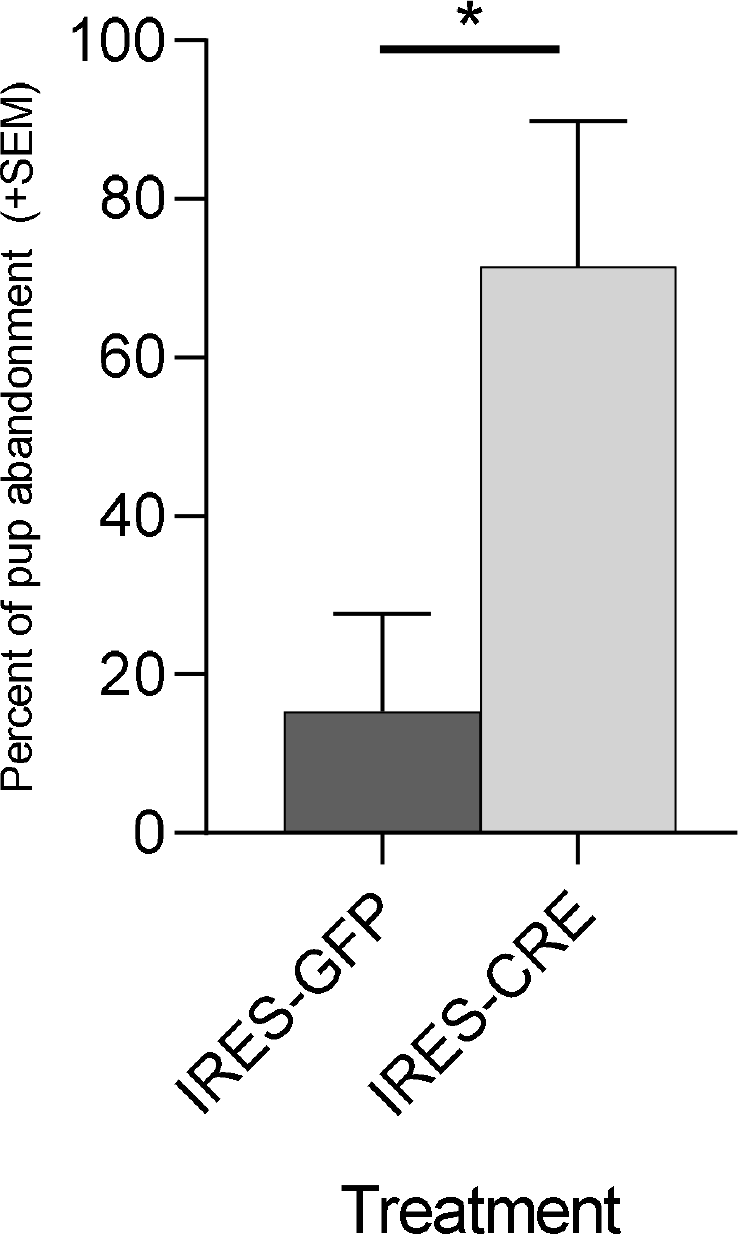
Percent of litters abandoned by Oxtr^flox/flox^ dams that received either control (IRES-GFP) or Cre (IRES-CRE) targeted injections in the nucleus accumbens (NAcc) shell. A significant genotypic difference in pup abandonment by treatment was observed. The females injected with Cre-IRES-GFP displayed increased abandonment by PND2 compared to dams injected with IRES-GFP. Graphs depict mean + SEM (*p≤0.05).

**Figure 12:**
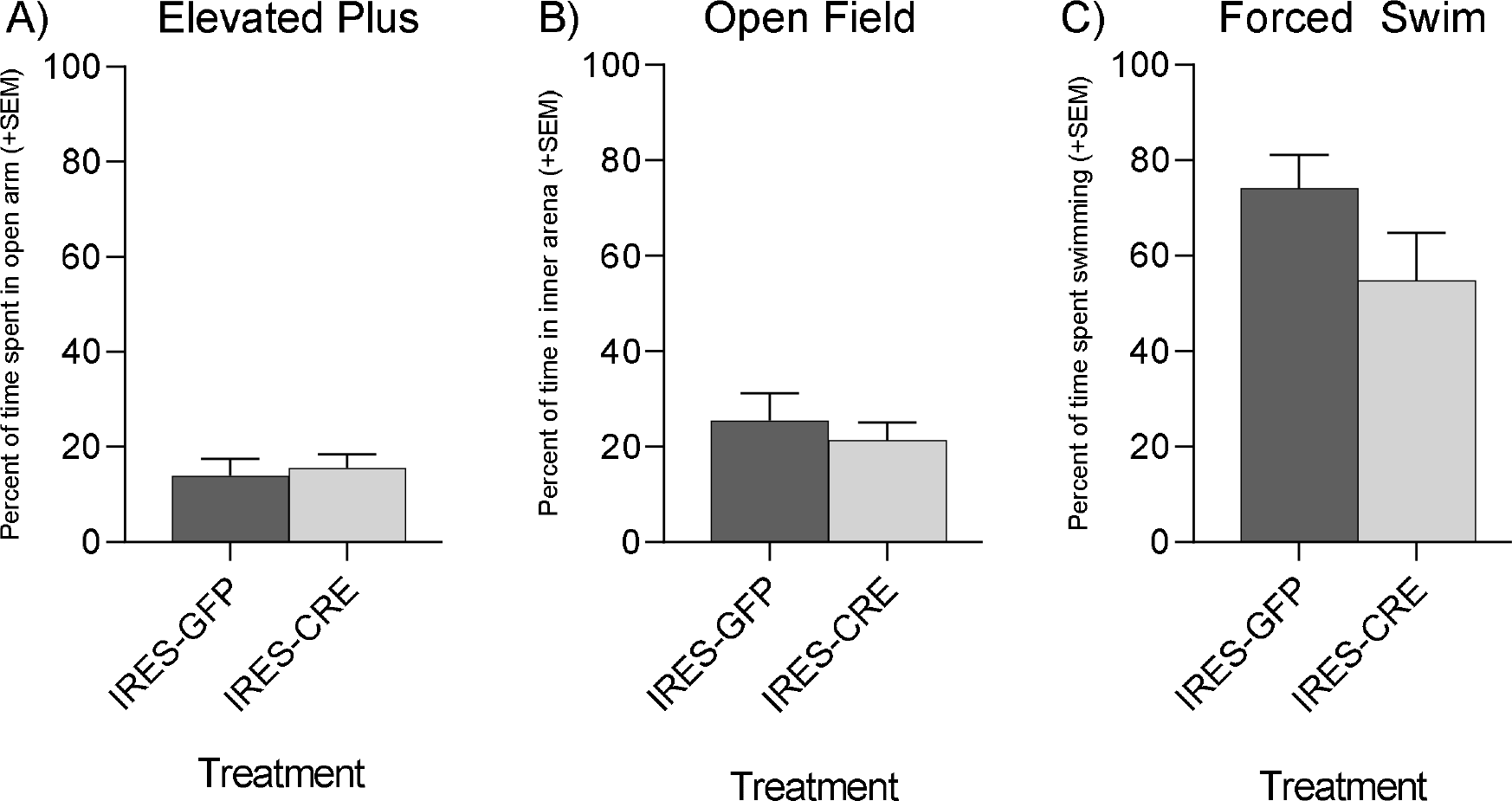
Anxiety- and depressive-like behaviors as measured by the A) elevated plus, B) open field task and C) forced swim in Oxtr^flox/flox^ dams that received either control (IRES-GFP) or Cre (IRES-CRE) targeted injections in the nucleus accumbens shell. All dams were tested four days after parturition regardless of pups presence in cage. No significant difference by treatment was observed between groups in percent time spent in the open arms of elevated plus, inner arena of open field or swimming in the forced swim task. Graphs depict mean + SEM.

## Discussion

Based on the findings of this work, we have provided further support for the idea that sensitized vs. maternal are not neurochemically the same. More importantly, we have identified a neuroanatomical substrate where, following parturition, Oxt signaling through Oxtr mediates the onset of maternal behavior in post-parturient rodents. Following consistent increases in c-Fos immunoreactivity within both one-hour post-parturient Oxtr −/− and Oxtr FB/FB dams, the NAcc shell was established as a region of interest. We further demonstrated the importance of Oxtr within the NAcc shell by genetically targeting and knocking out the Oxtr in this brain region, resulting in a pup abandonment phenotype similar to that observed in Oxtr −/− (24) and Oxtr FB/FB dams (25). Specifically, 71% of dams injected with CRE-IRES-GFP abandoned their pups.

In sensitized Oxtr FB/FB virgin females there were no observed differences in c-Fos immunoreactivity in the NAcc shell, as was observed in Oxtr FB/FB post-parturient dams. Rather, we found a difference in the LSD, with Oxtr FB/FB having lower activation than controls. Based on known differences between pup-sensitized females and post-parturient dams, this finding was not completely surprising (14). Further, the only behavioral differences in virgin Oxtr FB/FB females versus controls were on day one, with increased latency in retrieval of first pup observed in the Oxtr +/+ compared to Oxtr FB/FB. Thus, we speculate that increased c-Fos activation in the LSD of Oxtr +/+ virgin females may contribute to the innate avoidance of pups.

Decreases in social fear in dams is associated with decreases in the activation of the LS, but increases in Oxt release (39). The LS also regulates emotional processes and stress responses via dense interconnections with several limbic, diencephalic, and midbrain regions and has connectivity to the MBNN at the BNST, NAcc, PAG, and VTA (40). Oxtr expression is localized to the LS and there is evidence that Oxt in the LS can enhance social memory (41-43). Further, in lactating dams, increases in Oxt fiber density from the SON to the LS are critical for decreasing social fear (44). Based on these data, we postulate that activation of the Oxtr in the LSD increases as virgin females are continually exposed to the pups. This idea is supported by a recent study in which sensitized females, but not females that were exposed to pups one at a time, showed increases in neuronal activity in the mPOA and Oxt system (32). This idea could be tested by evaluating changes in c-Fos immunoreactivity across days of sensitization between the Oxtr +/+ and Oxtr FB/FB females.

Beyond day 1, there were no genotypic differences in maternal behaviors following sensitization, which is consistent with the previously reported normal maternal care in post-parturient Oxtr FB/FB dams. Certainly, sensitized females do differ from non-sensitized females. For instance, while laboratory mice display a spontaneous maternal behavior in which they retrieve and adopt a nursing posture over pups within minutes of exposure (45), only sensitized virgins will retrieve pups from a novel environment back to their home cages (46). As stated earlier, the observed differences in c-Fos activation in sensitized, as compared to post-parturient, females does highlight the importance of studying pre- and post-parturient females in order to understand the maternal brain, rather than sensitized females. However, it is the findings from Experiment 2 that support our *hypothesis* that the pup abandonment phenotype observed in Oxtr −/− and Oxtr FB/FB females (24, 25) is likely due to a failure of the brain to “switch” to a more maternal state. Further, our data from Experiment 3 suggest that this “switch” is located in the NAcc shell.

Genetic disruption of Oxtr expression in the NAcc shell resulting in a robust pup abandonment phenotype suggests that Oxtr expression in the NAcc shell is a critical for the onset of maternal behavior in females that have undergone parturition. Interestingly, Oxtr expression within the NAcc differs between species, with expression being generally higher in species that are more responsive to their young (47). In prairie voles, for instance, Oxtr expression in the NAcc has been established as being important for the neural regulation of alloparental care, i.e. parental care towards non-descendant young, and pair bond formation (34, 48-51). Though, the work described here is the first to functionally link Oxtr expression in the NAcc to maternal care in postparturient females.

The NAcc is part of the MBNN, which is important for maternal motivation and memory. In hormonally primed females, dopamine (DA) is released into the NAcc following activation of the VTA, via projections from the mPOA/vBNST. The NAcc receives also receives afferent projections from the BLA/PFC and DA is known to dampen the response to BLA/PFC input. Thus, when DA is present there is a dampening of NAcc output. The consequence of this is that the VP is released from GABAergic inhibition, this disinhibition is thought to allow the VP to be responsive to pup stimuli — which is permissive for appetitive maternal behaviors (52). In our current study, we were not evaluating the dampening of NAcc output, but we did observe an increase in immediate early gene activation in the NAcc of both Oxtr FB/FB or Oxtr −/− dams in measures of immediate early gene activation. Thus, we speculate that genetic disruption of Oxtr signaling resulted in the GABAergic ‘brake’ on the VP to be maintained, which would in turn inhibit the onset of maternal behavior. Still in question though, is where exactly the Oxt is coming from as well as the phenotype of the cell(s) on which the Oxtr is expressed.

### Conclusions

Taken together, the data presented here shed light on the importance of the Oxt system both in the maternal sensitization model and perhaps more importantly for the onset of maternal care in post-parturient females. Differential neuronal activation was observed depending on the presence, or absence (sensitization), of hormonal changes during parturition. Based on our findings from Experiment 1, we hypothesize that the observed decreases in maternal responsiveness, as measured by pup retrievals, on Day 1 in Oxtr FB/FB sensitized virgins may be due to the loss of Oxtr inhibition within the LSD, ultimately altering the virgin’s response to pup cues. In Experiment 2, the NAcc shell was identified as a neuroanatomical region where Oxt signaling via the Oxtr may play a role in the onset of maternal care. This was confirmed in Experiment 3 when the targeted knockdown of Oxtr in the NAcc shell resulted in a pup abandonment phenotype similar to what has previously been observed in Oxtr FB/FB and Oxtr −/− dams. Looking forward, additional studies will need to be conducted to determine which aspects of pup cues the Oxt system is modifying. For instance, it would be interesting to know if the loss of Oxtr within the NAcc shell altered the dam’s motivation/attraction to pups. Given the relative conservation of the Oxt system across species, it is likely that the role of the Oxt system in the onset of maternal behavior is conserved in other mammalian species. Thus, this work has implications beyond just mice.

## Materials and Methods

### Animals

Adult females in this study were generated from three transgenic mouse lines: Oxtr −/−, Oxtr FB/FB, and Oxtr flox/flox mice (35, 36). The Oxtr −/− and Oxtr FB/FB mice were generated from heterozygous breeding pairs with Oxtr +/+ siblings as controls and the Oxtr^flox/flox^ females from homozygous lines. All mice were bred in the Kent State University vivarium and maintained on a 12:12 light/dark cycle, with food and water provided ad libitum, except during behavioral testing. All animals were weaned 21 days postpartum and housed in single-sex sibling groups. Tails were clipped at the time of weaning, DNA extracted and genotyping conducted as previously described (35). All experiments were conducted in accordance with the Kent State University Animal Care and Use Committee.

### Experiment 1: Maternal sensitization and immediate early gene activation in Oxtr +/+ and Oxtr FB/FB dams

#### Experiment 1a: Pup Sensitization

Adult female Oxtr +/+ and Oxtr FB/FB mice were single housed one week prior to testing. Each female was exposed to four newly born C57BL/6J pups (two males and two females) for 30 minutes for four consecutive days (postnatal day (PND) 1-4); we used this approach to avoid any confound associated with having litters of mixed genotype. Exposure started once the first pup was placed in the cage and only females who retrieved all four pups within the first five minutes continued to have their maternal behavior scored. If females failed to retrieve the pups within the first five minutes, all pups remained in the cage for the full 30 minutes to ensure sensitization. On PND4, one hour after the last exposure, females were euthanized, and their brains removed and placed into fixative (4% paraformaldehyde); in preparation for c-Fos immunocytochemistry.

#### Maternal behavior scoring

The behavior of sensitized females was scored by an observer with no knowledge of genotype/treatment using Observer 5.0 (Noldus, Leesburg, VA). Sensitized females were scored for 1) pup interactions (sniffing/licking); 2) nonsocial behaviors (resting alone/feeding); 3) nest building; and 4) self-grooming. To evaluate potential anxiety-like behaviors, we further broke down nonsocial behaviors into digging, rearing, and exploratory behaviors. Pup retrieval latencies were measured by determining the amount of time it took the female to retrieve the first and last pups. Maternal behaviors, except for pup retrievals, were analyzed within each day using a two-way ANOVA with genotype as the main factor. If a p value of ≤ 0.05 was found, a Fisher’s least significant differences *post hoc* test was performed. For this analysis, the amount of time the sensitized females engaged in all behaviors (maternal and nonmaternal) were first summed, and the percentage of time the females engaged in each behavior was calculated. Females had five minutes to retrieve the pups, if they did not retrieve the pups then no maternal behavior was scored for that day and a 300 second latency score was assigned. The latency to retrieve the pups was analyzed within each day using a one-way ANOVA. A p value of ≤ 0.05 was considered statistically significant.

#### Experiment 1b: c-Fos immunocytochemistry and quantification

Fixed brains were cut into three series of 50μm free-floating sections using a Vibratome 1000Plus (Leica Microsystems, Buffalo Grove, IL) and stored at −20°C in cryoprotectant (0.1M potassium phosphate buffer, sucrose, polyvinyl pyrolidione and ethylene glycol) prior to staining. At the time of staining, sections were washed six times for five min in 1XPBS, incubated in 1.5% hydrogen peroxide for five min, and then washed two times for five min. in 1XPBS. Following the washes, the sections were incubated in 1XPowerblock™ (Universal Blocking Reagent 10X, BioGenex, Fremont, CA) for 30 min. After blocking, sections were moved into the primary antibody (Santa Cruz Biochemicals, Santa Cruz, CA, USA, rabbit anti-c-Fos, sc-52) at 1:5000 in antisera diluent (PBS + 1% normal goat serum + 0.3% Triton X-100) and incubated overnight at 4°C. The next day the sections were washed in 1XPBS three times for five min. and then incubated in avidin-biotin complex (Vectastain Elite ABC (Rabbit IgG), Vector Laboratories (Burlingame, CA, USA, PK-4001)) for one hour at room temperature. Sections were then washed in 1XPBS three times for five min. and incubated with diaminobenzidine (DAB) for 2–10 min. (Vector Laboratories, Burlingame, CA, USA, DAB substrate kit, SK-4100). To deactivate DAB, the sections were rinsed with 1XPBS followed by two washes in 1XPBS five min. The tissue was sequentially organized and mounted onto gel-subbed slides, allowed to air dry and cover slipped using DPX Mounting Medium (Sigma-Aldrich, DPX 06552).

#### Quantification

c-Fos immunoreactive cells were counted at 100X magnification by an observer with no knowledge of the testing groups. iVision software (BioVision Technologies, Exton, PA) was used to capture the images and c-Fos-ir cells were manually counted within each neuroanatomical area. Three sections per area, with sections being 100μm apart, were bilaterally counted and averaged using set box sizes for each area (box sizes from (37)), which served to normalize count area across animals. The areas measured included the dorsal and ventral aspects of the lateral septum, the LSD (732 x 754 pixels) and the LSV (380 x 338 pixels), respectively, the dorsal and ventral aspects of the BNST, the BNSTD (600 x 870 pixels) and the BNSTV (600 x 435 pixels), respectively, the mPOA (465 x 870 pixels), VTA (300 x 300), MeA (990 x 870), periaqueductal grey (PAG) (300 x 550), lateral habenula brain region (LHB) (300 x 300) and NAcc core (500 x 500) and shell (500 x 500). These areas were identified based upon Paxinos and Franklin mouse brain atlas. Comparisons between cell counts were made by area between genotypes using a one-way analysis of variance (ANOVA) (SPSS 16.0 for Mac, IBM, Armonk, NY). A result was considered statistically significant if p ≤ 0.05.

### Experiment 2: Parturition-induced immediate early gene activation in Oxtr +/+, Oxtr −/−, and Oxtr FB/FB dams

#### Breeding

Females were group housed 2-4 per cage for two weeks. At the end of this two weeks, male bedding was added to female cages one day prior to the introduction of males to induce estrous (38). The following day C57BL/6J sires from our colony were placed into the female grouped-housed cages. Each day, for one week following pairing, females were checked each morning for the presence of a sperm plug to indicate a possible pregnancy. If a sperm plug was found, the male was removed, and the female was single housed. At the end of one week, all remaining males were removed from the females’ cages. Females that were single housed were then monitored for pregnancy based on weight gain. Any females, who were not pregnant, were paired again after two weeks had passed. Starting on gestational day 18, dams were checked hourly so that we could “catch” parturition. When a dam gave birth, she was euthanized by cervical dislocation one hour after the birth of the last pup, confirmed by gentle palpation of abdomen. Brains were collected and stored in 4% paraformaldehyde until processed for c-Fos immunocytochemistry as described for Experiment 1.

### Experiment 3: Effects of targeted genetic knockdown of the Oxtr on maternal behavior

Based on the findings of Experiment 2, we evaluated whether or not disruption of Oxtr signaling in the NAcc shell resulted in any quantifiable changes in either the onset or expression of maternal behavior.

#### Intracranial Injections

Prior to surgery, 16 Oxtr flox/flox sibling females were randomly assigned to two groups: 1) Cre recombinase adeno associated virus (Cre-IRES-GFP) (n=8) or 2) control (IRES-GFP) (n=8). The Cre-IRES-GFP (AAV2/2CMVCRE-wtIRESeGFP) and IRES-GFP (AAV2/2CMVeGFP), were purchased from the University of Iowa Viral Vector Core. At the time of surgery, animals were anesthetized using 2% isoflurane/oxygen mixture and placed into an Ultraprecise stereotaxic apparatus (David Kopf Instruments, Tujunga, CA). Once animals were secured in the ear bars, a midline incision was made across the top of the skull. The injection target was the NAcc shell and the coordinates used were, from bregma: anterior posterior +1.58, medial lateral ±0.7 and dorsal ventral −4.5 (from top of the skull). Burr holes were made using a hand-held drill (Dremel®, Racine, WI) with an engraving cutter bit (model #105, Dremel®, Racine, WI). Once the burr hole was created, the needle of a 2µL Hamilton syringe (Hamilton Company, Reno, NV) was placed at the appropriate depth and allowed to sit for five minutes prior to injection to allow the brain to reposition. Bilateral injections of 0.5μl of either the Cre-IRES-GFP or IRES-GFP were injected at a rate of 0.2µl/min to limit damage. The needle remained in place an additional five minutes to allow for diffusion of the virus. The needle was then slowly removed, and the skin brought back together over the skull and closed with a wound clip. Following surgery, animals were administered 0.3 ml of warm saline (0.9%) intraperitoneally (i.p.) to aid in recovery and single housed with microisolator lids. Enrichment provided included a red dome and nestlets plus dams received peanuts at the time of cage changes. All animals were given two weeks to recover, to allow for maximal viral infection and the down regulation of Oxtr before being paired with a C57BL/6J sire from our colony.

#### Mating

Females that underwent stereotaxic surgery were paired with C57BL/6J sires from our colony. All females were weighed weekly and once females appeared pregnant (or gained >3g), males were removed from their cages. Two weeks after the male was removed females were re-paired if they did not appear pregnant. Additionally, if a female’s first litter did not survive, she was re-paired to evaluate the effects of the loss of the Oxtr within the NAcc shell on the care of subsequent litters.

#### Pup Observations

Dams were checked twice daily (0900h and 1400h), PND0 was designated as first day pups were present in the cage and completion of birth was confirmed via gentle palpation of dam’s abdominal section. A general health check of dams and pups occurred following parturition, including presence of a milk spot, number of pups (alive and dead), sex of pups and group pup weights. All litters were culled to 4 pups on PND0, 2 males and 2 females. To assure maternal behavior testing did not affect pups, the pup weights and general health checks were collected daily.

#### Maternal behavior

The first measure of maternal behavior was the evaluation of pup abandonment. Any experimental animals that did not abandon their pups were then tested for maternal behavior from PND1 to PND3. For the measures of maternal behavior, the animal numbers were n=8 for IRES-GFP and n=2 for IRES-CRE. Prior to all behavioral testing, animals were acclimated to the testing space for one hour after lights out under dim red-light illumination. Pup retrieval was evaluated by first removing all pups from the home cage for five minutes while the dam’s behavior was videotaped. Pups were then scattered opposite to the location of the dam in the home cage and the five-minute pup retrieval task was videotaped. Pup retrievals were later quantified and included the latency to retrieve the first pup and the latency to retrieve all pups. If all pups were not retrieved within five minutes of being returned to the cage, a latency score of 300s was recorded and no maternal behaviors were scored. Following the five minutes of retrieval the dam’s behavior was videotaped for an additional 20 minutes. If all pups were retrieved, 20 minutes of maternal behavior was scored. Behaviors scored included, time on/off nest, licking/sniffing pups, nursing/crouching nest building, self-grooming, rearing. Behavior was scored by an observer blind to genotype/treatment using Observer 5.0 (Noldus, Leesburg, VA) as previously noted.

#### Postpartum behaviors

##### Elevated Plus

On PND 4, experimental animals were tested on the elevated plus maze as previously described (24). Briefly, all animals were moved to the room one-hour prior to testing and the maze was illuminated at approximately 100 lux. Animals were placed in the center facing the closed arms of the elevated plus maze and electronically tracked for 10 minutes using EthoVision XT (Noldus, Leesburg,VA). The elevated plus maze consists of three quadrants: open arms (10 cm x 45 cm), closed arms (10 cm x 45 cm x 40 cm), and the center platform (10cm x 10 cm). To assess anxiety-like behavior between treatment groups, the duration of time spent in the open and closed arms was summed and the percentage of time spent in the open and closed arms was determined.

##### Open Field

On PND 5 experimental animals were tested in an open field test, as previously reported (24). Animals were moved to the testing room one hour prior to testing, the arena was illuminated at approximately 200 lux. All animals were tested in the open field made out of Plexiglas measuring 45.5×45.5×30 cm. Animals were placed into the center of the open field and movement tracked for a total of 20-minutes by Ethovision XT (Noldus, Leesburg, VA). For analysis of anxiety-like behavior, the field was separated into two parts, an inner arena (measuring 32×32 cm) and outer arena. The tracking system quantified the amount of time spent in the inner versus outer arena. From this, the percentage of time spent in the inner and outer arenas was calculated.

##### Forced Swim

The forced swim test was administered on last day of testing (PND 6) to assess depressive-like behaviors. Animals were moved to the testing room one-hour prior to testing, which occurred during the dark phase. Animals were placed into a 19cm diameter cylindrical tank which was ¾ full of room temperature (∼21°C) water. Animals were videotaped for six minutes and for the duration of the test they were observed for any signs of distress. All dams were then returned to a clean cage to which their nest and pups had been moved. Forced swim videos were later scored using Observer XT 9 (Noldus, Leesburg,VA) for swimming or floating for a total of four minutes, beginning at minute two, to allow for acclimation. To reduce observer error associated with transition, we used a sampling method of scoring. Behavior was scored every five seconds as either swimming behavior, two or more paws moving to propel the mouse, or float behavior consisting of no paws moving or two or fewer paws moving slightly only to stabilize the mouse in the water. The number of float and swim behaviors scored were summed and the percentage of swim and float behaviors scored was determined. All postpartum behavioral measurements were compared between genotypes using a one-way ANOVA with treatment groups as the main factor. A p-value of ≤0.05 was considered statistically significant.

##### Site Checks

Following behavioral testing, intracranial injected animals were euthanized, brains fast frozen and stored at −80°C for injection site confirmation using green fluorescent protein (GFP) immunostaining. Tissue was sectioned at 12µm in a −20°C cryostat (Leica 1950; Leica Biosystems, Buffalo Grove, IL, USA) and mounted onto Superfrost Plus slides (Fisher Scientific, Hampton, NH, USA). On the first day of staining, sections were fixed for five min using 4% paraformaldehyde at room temperature and rinsed four times with 1XPBS prior to five minutes was in 1XPBS. Sections were placed in 1X Power block for 10 minutes (Universal Blocking Reagent 10X, Biogenex, Cat#HK085-5K), repeated rinse and wash with 1XPBS and placed in rabbit anti-GFP primary (1:20,000 in 1%BSA, 1XPBS) overnight at 4°C. The next day, section were rinsed with 1XPBS then washed three times for three minutes each. Sections were placed into 1.5% H_2_O_2_ for 20 minutes at RT, washed for four times, three minutes each with 1XPBS and gently dried. Super Picture HRP Polymer Conjugate Rabbit Primary Kit (Invitrogen, Cat#87-9263) was used for the following steps. Each slide had 100µl of antibody from Rabbit PolyHRP conjugate applied and incubated for 30 minutes at RT. Washed two times in 1XPBS for three minutes each and one time in 0.1M Tris (pH8) for three minutes. The DAB step was prepared according to Rabbit PolyHRP directions. Sections were air dried and cover slipped using DPX mounting media. Site checks were then performed with light microscopy.

